## Supplemental Information for "Cortical Plasticity is associated with Blood-Brain-Barrier Modulation"

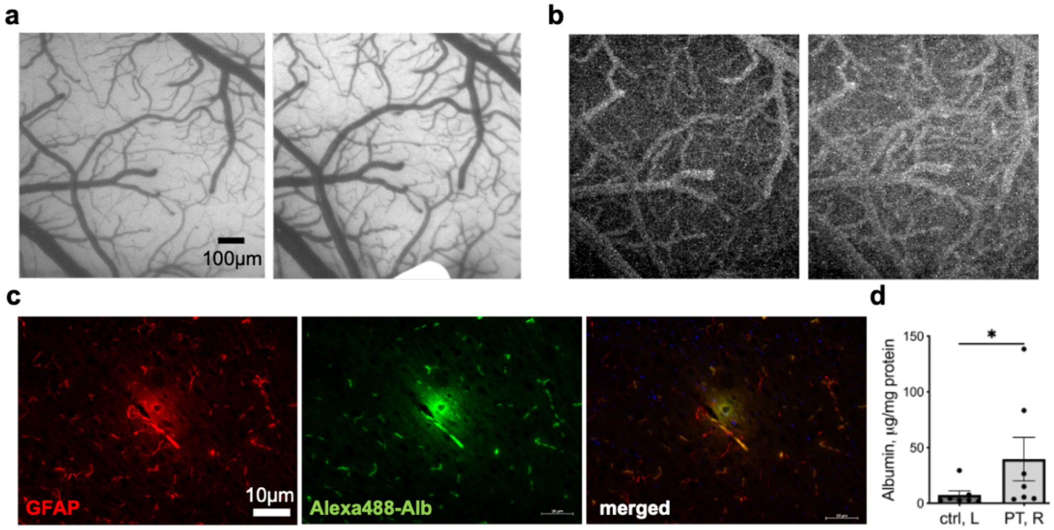


**Supplemental Figure S1: Increased BBB permeability following stimulation. a.** Images of the responding region of the rats' cortex before (left) and after stimulation (right). **b.** Images depicted in (a) after NaFlu injection before (left) and after (right) stimulation. **c.** Representative image of a small vessel in the stimulated hemisphere stained for astrocytes (GFAP, left) albumin (middle) and the merged image (right) showing co-localization of the two markers and albumin accumulation surrounding the vessel. **d.** Albumin concentration in the contralateral hemisphere relative to the ipsilateral in rats following PT-induced BBB dysfunction. *$p<0.05$, Wilcoxon, n=7, mean ± SEM.

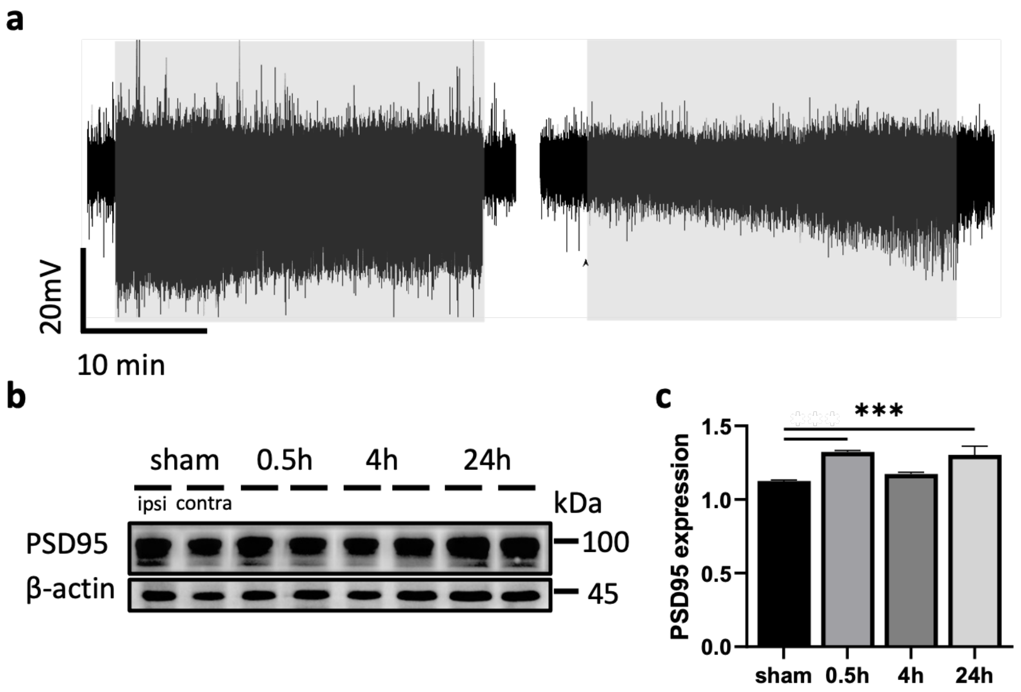


**Supplemental Figure S2: Modulation of BBB permeability associated with synaptic plasticity is activity dependent. a.** Representative LFP trace from the responding area in L2/3 of the somatosensory cortex during a 30 min stimulation following AP5 (left) or CNQX application (right); Greyed rectangles mark stimulation. **b.** Representative western blot for PSD-95 expression in the ipsi- and contra-lateral cortex of sham and stimulated rats at different time points. Normalized to β–actin (loading control). **c.** Quantification of PSD-95 expression levels at different time points post stimulation. (n=8 each group, one-way ANOVA with FDR correction, ***p<0.001).

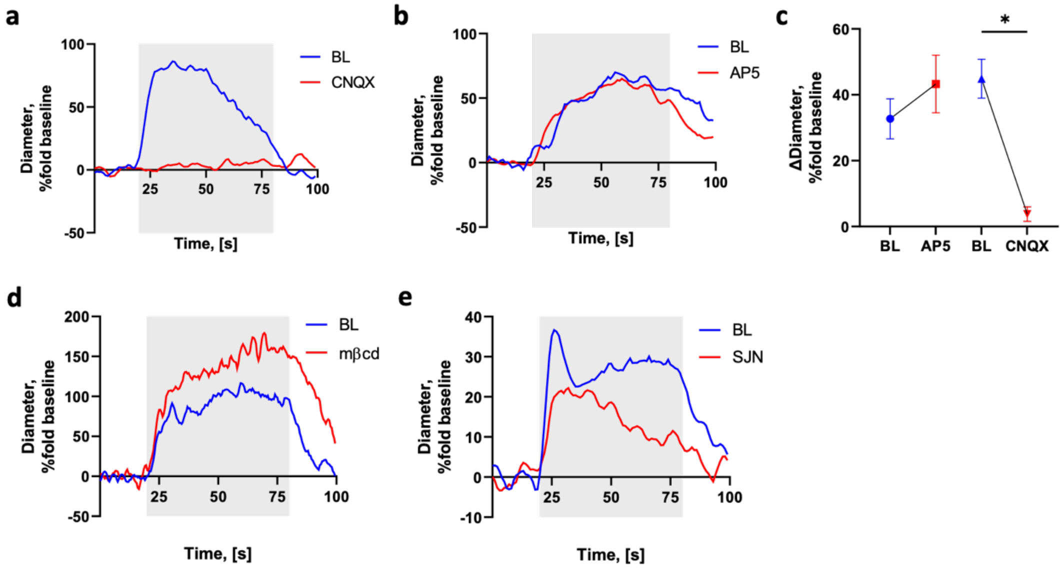


**Supplemental Figure S3: Stimulation-induced BBB modulation can be prevented with or without affecting the vascular response. a-b.** Representative examples of the responding arteriole diameter during 1 min stimulation. Presented as % change from a 20 s baseline segment before stimulation. Blue line, baseline (BL); red line, following application of AP5 (a) or CNQX (b). **c.** Mean arteriolar diameter during 1 min baseline is similar compared to AP5 and significantly larger compared CNQX stimulations. Presented as % change from a 20 s baseline segment before stimulation (AP5, n=2; CNQX, n=3; p=0.011 paired t-test). **d-e.** Representative examples of the responding arteriole diameter during 1min stimulation, presented as % change from a 20 s baseline segment before stimulation. Blue line, baseline (BL); red line, following application of mβCD (d), SJN (e).

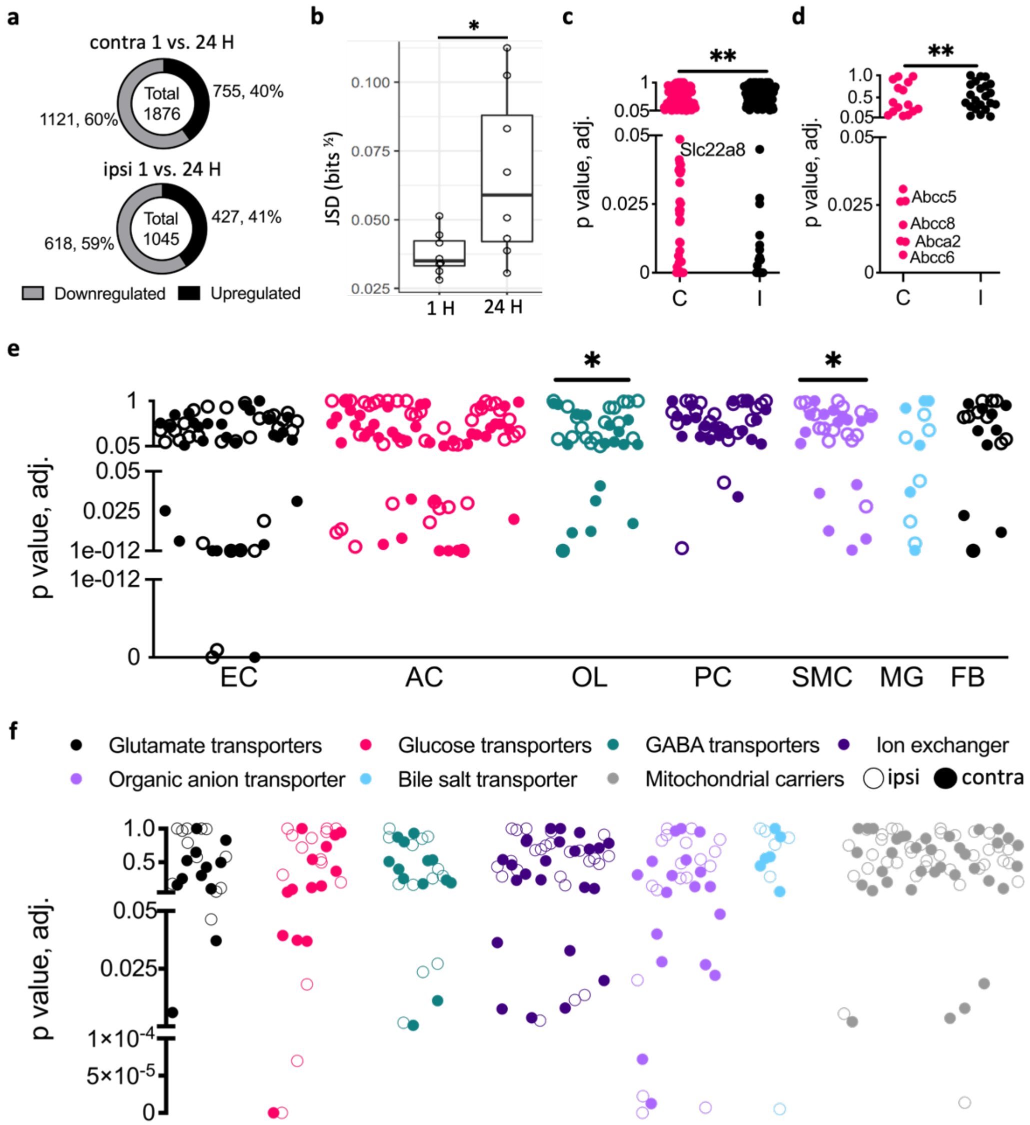


**Supplemental Figure S4: Neuronal activity regulates BBB transport and NVU cells-specific genes. a.** Significant differentially expressed genes (DEG) in the contralateral cortex 24 vs. 1 h after stimulation (top) and same for the ipsilateral cortex (bottom). **b.** Jensen–Shannon divergence (JSD) calculations using normalized counts indicate statistically significant difference between paired contra and ipsi cortical RNA expression of the rats 24 vs. 1 h after stimulation (n=8 each, mean ± SD, p=0.034). **c-d.** Scatter plots indicating solute carrier transporters (Slc, **c**) and ATP-binding cassette transporters (ABC, **d**) gene expression in the contra vs. ipsi cortices. (C-contra, I–ipsi, Slc n=331, p=0.0069, ABC n=45, p=0.0059, Chi-square). **e.** Scatter plot of NVU cell specific DEGs in the contra vs. ipsi cortices of stimulated rats. (OL n=19, p=0.0182, SMC n=16, p=0.035, Chi-square **f.** Scatter plots of Slc transporter genes grouped by families from the ipsi (empty circles) and contra (filled circles) cortices.

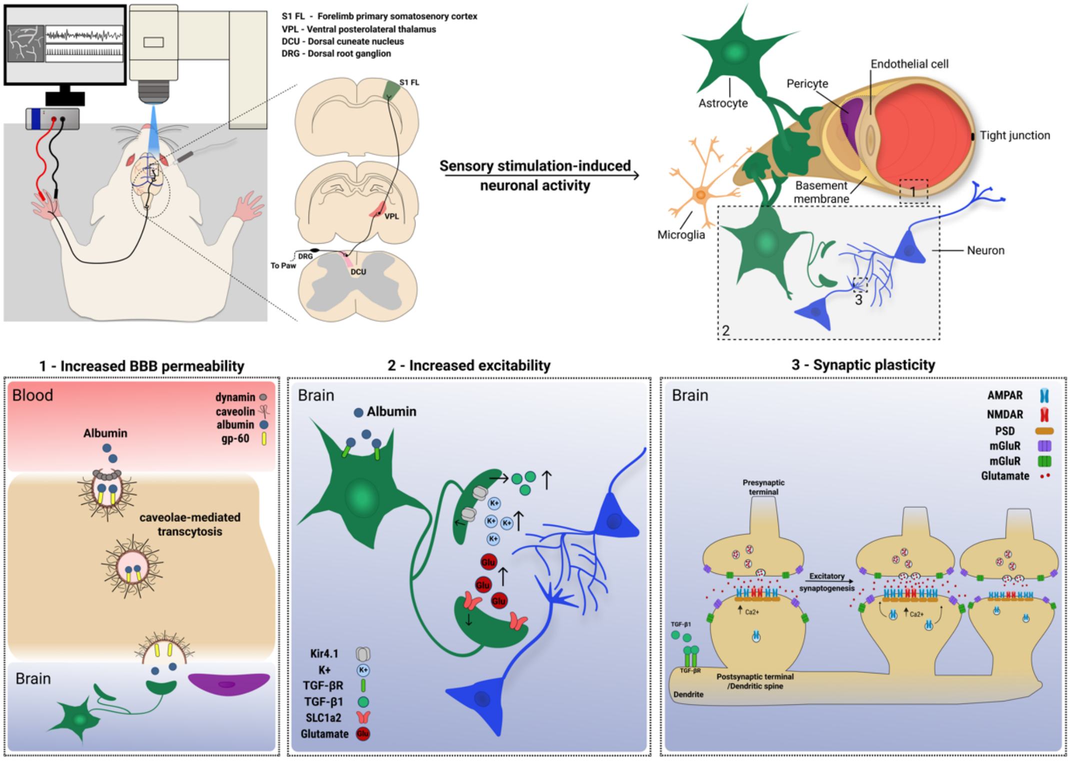


**Supplemental Figure S5: Activity-dependent modulation of BBB permeability associated with synaptic plasticity.** Top left: Experimental procedure illustrating stimulation of the forepaw to elicit neuronal activity in the somatosensory cortex of the rat. Top right: Illustration of the NVU. Labeled rectangles indicate the hypothesized processes induced by the stimulation. Bottom: 1. Stimulation-induced increased BBB permeability manifested by increased CMT of serum albumin. 2. Increased excitability is induced by albumin binding to TGF-β receptors on astrocytes (Weissberg et al., 2015), leading to secretion of TGF-β1(Allen & Eroglu, 2017), and downregulation of Kir4.1 channels and glutamate transporters (David et al., 2009; Frigerio et al., 2012), which in turn result in elevated extracellular $K^{+}$ and glutamate (David et al., 2009; Ivens et al., 2007). 3. Increased excitability leads to synaptic plasticity, indicated by excitatory synaptogenesis (Patel & Weaver, 2021).
